# Non-redundant functions of H2A.Z.1 and H2A.Z.2 in chromosome segregation and cell cycle progression

**DOI:** 10.1101/2020.09.15.297671

**Authors:** Raquel Sales-Gil, Ines J. de Castro, Hasnat Amin, Christos Spanos, Veronica Vinciotti, Juri Rappsilber, Cristina Sisu, Paola Vagnarelli

**Author notes:** Correspondence and requests for materials should be addressed to P.V.

## Abstract

H2A.Z is a H2A-type histone variant essential for many aspects of cell biology ranging from gene expression to genome stability. From deuterostomes, H2A.Z evolved into two paralogues H2A.Z.1 and H2A.Z.2 that differ by only three amino acids and are encoded by different genes (*H2AFZ* and *H2AFV* respectively). Despite the importance of this histone variant in development and cellular homeostasis, very little is known about the individual functions of each paralogue in mammals. Here we investigated the distinct roles of two paralogues in cell cycle regulation. Using a specific siRNA approach for each paralogue in human cells, we unveiled non-redundant functions for H2A.Z.1 and H2A.Z.2 in cell division: H2A.Z.1 regulates the expression of important cell cycle genes (including Myc and Ki-67) and its depletion leads to a G1 arrest, whereas H2A.Z.2 is essential for centromere integrity and function, thus playing a key role in chromosome segregation.

## INTRODUCTION

Nucleosomes form the basic unit of eukaryotic chromatin and consist of 146 DNA base pairs wrapped around an octamer of histone proteins. Canonical histones are incorporated into nucleosomes during DNA replication but histone variants, encoded by separate genes, are typically incorporated throughout the cell cycle ^1-3^.

H2A is one of the four core histones. Sequence analyses have shown large-scale divergence in the H2A family, resulting in numerous variants ^4^. H2A.Z, originally identified in mouse cells ^5^, is a highly conserved H2A-type variant. H2A.Z is present as a single variant until early deuterostomes when two H2A.Z paralogues appear: H2A.Z.1 and H2A.Z.2. These variants differ by only three amino acids and are encoded by the *H2AFZ* and *H2AFV* genes, respectively ^6^. More pronounced differences are found in their regulatory regions. Specifically, the promoter sequence of H2A.Z.1 is similar to the one of other histone variants (it contains a TATA box, three CAAT boxes and several putative GC-boxes), whereas H2A.Z.2 promoter lacks a TATA box and the CAAT regions ^7^. In addition, H2A.Z.1 and H2A.Z.2 present a different L1 loop structure and studies using fluorescence recovery after photobleaching (FRAP) showed that H2A.Z.2-containing nucleosomes are more stable than the H2A.Z.1-containing ones ^8^. In primates, H2A.Z.2 has two splice variants: H2A.Z.2.1 and H2A.Z.2.2, where H2A.Z.2.2 has a shorter docking domain and forms highly unstable nucleosomes ^9^.

Although H2A.Z knockdown leads to early embryonic lethality in Drosophila ^10^ and mice ^11^, depletion of the H2A.Z orthologue in *S. cerevisiae*, HTZ1, is not lethal ^12^, pointing out to a possible difference on the role of H2A.Z among species.

Several studies have highlighted the importance of H2A.Z in transcription regulation ^13-17^. However, whether H2A.Z promotes or represses transcription appears to depend on the gene, chromatin complex, and post-translational modifications of H2A.Z itself ^18^. In several organisms, H2A.Z peaks at transcriptional start sites (TSS) of active and repressed genes ^19-22^ and specifically localises at the +1 nucleosomes in the direction of transcription ^23^.

H2A.Z is also linked to heterochromatin regulation. Recent evidence suggests that H2A.Z and H3K9me3, a known marker for heterochromatin, can cooperate to enhance the binding of Heterochromatin Protein 1 alpha (HP1α) to chromatin *in vitro* ^22 24 25^.

However, the specific contribution of each paralogue towards these quite different aspects of chromatin biology is currently unknown. Although H2A.Z.1 and H2A.Z.2 are distributed similarly around the nucleus and are subjected to comparable post-translational modifications, their 3D structure, genome localisation and tissue distribution appears to be quite different ^7, 8^. In fact, H2A.Z.2 does not compensate for the loss of H2A.Z.1 *in vivo*, as H2A.Z.1 knock-out is lethal in mouse ^11^.

To date, very few studies have attempted to investigate and differentiate the specific roles of these two paralogues in vertebrates. In DT40 chicken cells, knock-out of H2A.Z.2 results in a slower cell proliferation rate compared to the wild type and H2A.Z.1 knock-out cells ^26^, while in humans, Floating-Harbor syndrome ^27^ and malignant melanoma ^28^ have been specifically linked to H2A.Z.2 ^27^. In addition, both variants seem to play independent roles in the transcription of some genes involved in the response to neuronal activity ^29^. Moreover, very recently, it was shown that the differences between H2A.Z.1 and H2A.Z.2 on transcription regulation seems to depend more on the relative level of the two paralogues rather than on their chromatin localisation ^30^. However, we are still missing a full understanding of the role of each variant in human cells.

As the role of histone variants in genome organisation and regulation becomes more appreciated and emerging studies have linked H2A.Z to cancer, it is important to investigate and clarify the possible differential roles of the H2A.Z paralogues and splice variants for cell cycle regulation *in vivo*. This will not only provide a better understanding of their function, but it will also clarify their relative contribution to divergent aspects of chromatin biology. In this study we used siRNA to specifically knockdown H2A.Z.1 or H2A.Z.2 in human cells. Our results show for the first time that H2A.Z.1 and H2A.Z.2 perform non-redundant roles during cell cycle and chromatin organisation; whereas H2A.Z.1 is a key regulator of cell cycle progression and nuclear morphology, H2A.Z.2 controls chromosome segregation and heterochromatin regulation.

## RESULTS

### H2A.Z.2, but not H2A.Z.1, is necessary for heterochromatin maintenance in human cells

Several studies have linked H2A.Z to heterochromatin in different systems but it is still unclear if a subset of histone variants is particularly enriched at these genomic regions.

In order to study the chromatin flavour of HP1α-bound nucleosomes in an unbiased manner, we used a TAG-Proteogenomic approach recently developed in the lab ^31^ (Supplementary Figure 1 A). Briefly, HeLa nucleosomes prepared from SILAC-light labelled cells were incubated with recombinant GST:HP1α or GST alone. After elution and SDS-PAGE, the histone fractions were analysed by mass spectrometry. The SILAC heavy-labelled nucleosome fraction was used to normalise the abundance of the different histones before the binding to the GST column (Figure 1 A). Next, we analysed the abundance of different histone variants present in the GST:HP1α-bound fraction and compared it to the input (I) and GST alone. We used another chromatin-associated protein, SAP18 (which belongs to the Sin3A HDAC complex ^32^), as additional control (Supplementary Figure 1 C). The data revealed that H2A.Z is one of the histone variants specifically enriched in the HP1α-bound chromatin (Figure 1 B; Supplementary Figure 1 B, C).

**Figure 1.**
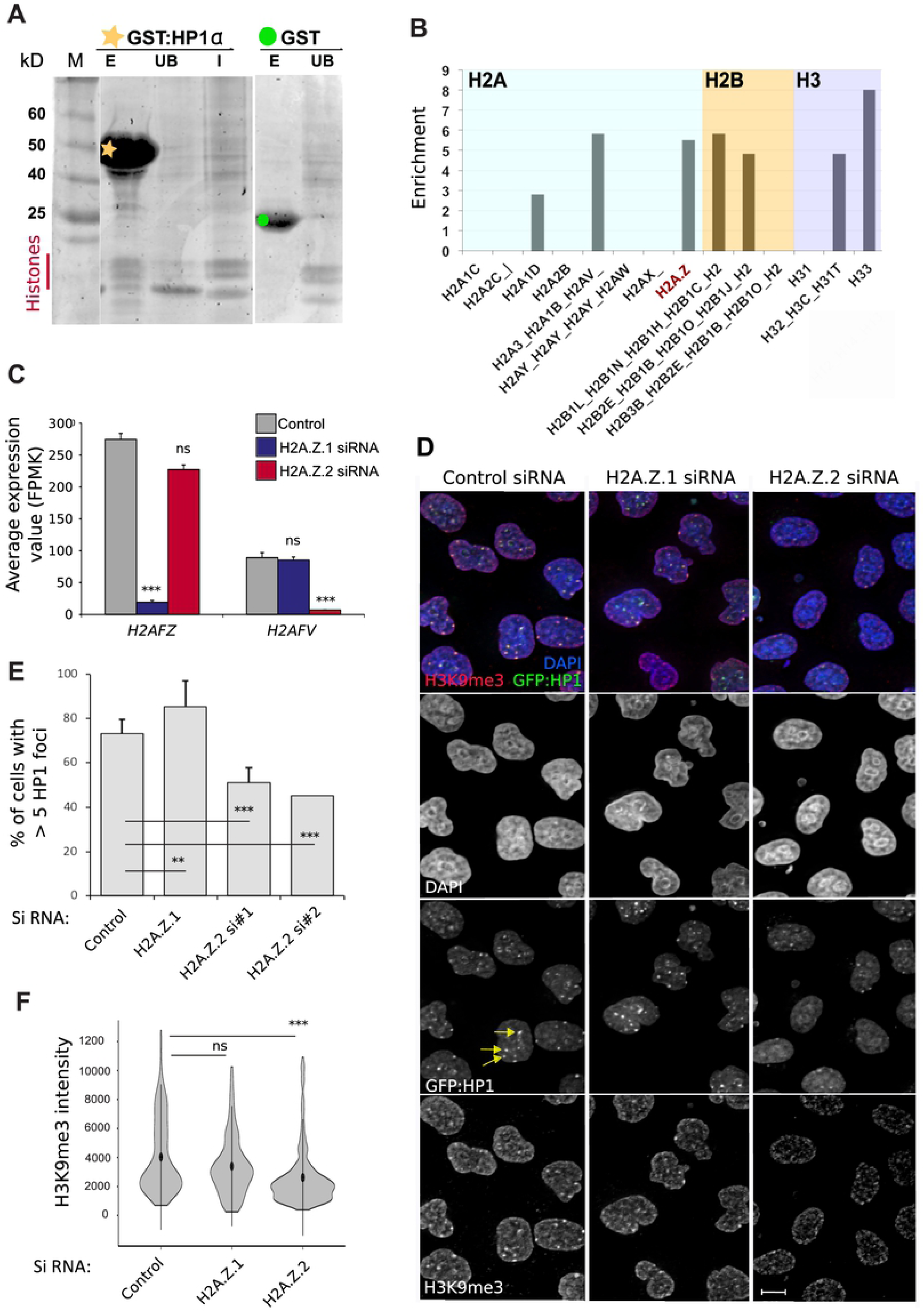
H2A.Z.2, but not H2A.Z.1, regulates heterochromatin. **A)** Recombinant GST or GST:HP1α were incubated with HeLa nucleosomes labelled with light amino acids (see procedure in De Castro et al., 2017). The calibration input (I) was labelled with heavy amino acids. The gel was stained with InstantBlue Coomassie Protein Stain and imaged by LI∼COR. The histones bands were excised from the gels and analysed by mass spectrometry. M= marker E= glutathione elution; UB= unbound fraction. **B)** Graph showing the enrichment of GST:HP1α pulled down-histones over GST after calibration against the input fraction. **C)** *H2AFZ* and *H2AFV* average expression values obtained by RNA sequencing of three biological replicates after control, H2A.Z.1 and H2A.Z.2 siRNA treatment. Error bars show the standard deviation (SD). ***=p<0.001; ns=not significant. **D)** HeLa GFP:HP1α (green) cells were transfected with control, H2A.Z.1 or H2A.Z.2 siRNAs for 72 h and stained for H3K9me3 (red). Yellow arrows indicate HP1 foci. Scale bar: 10µm. **E)** Quantification of interphase HP1α foci from experiment in (D). Error bars indicate SD of four biological replicates, except for H2A.Z.2_2 siRNA, where only one replicate was analysed. Data sets were statistically analysed using Chi-square test. At least 200 nuclei were analysed for each condition, except for H2A.Z.2_2 were 75 nuclei were analysed. **=p<0.01; ***=p<0.001. **F)** Violin plots representing the distribution of H3K9me3 intensity levels from experiment in (D). Mean with SD are shown. At least 170 nuclei, from three biological replicates were analysed for each condition. Data sets were statistically analysed using the Wilcoxon rank test in R. ***=p<0.001; ns=not significant. See also Supplementary Figure 1.

However, the mass spectrometry analysis alone did not allow us to distinguish between the two H2A.Z paralogues (H2A.Z.1 or H2A.Z.2). To investigate which H2A.Z isoform (if any) was important for HP1α maintenance *in vivo*, we used a HeLa GFP:HP1α cell line and specifically knockdown each H2A.Z variant by siRNA. We used two oligos designed against each variant and performed RNA-seq on the control, H2A.Z.1 and H2A.Z.2 siRNA-treated cells to confirm the specificity of the depletion. The data show that: a) the siRNAs are specific for each paralogue; b) in HeLa cells, as in many other systems, H2A.Z.1 is expressed at a much higher level than H2A.Z.2; c) the removal of one form does not interfere with the expression level of the other (Figure 1 C) or histone H2A (Supplementary Figure 1 D). Since there are no antibodies that can specifically distinguish between the two isoforms, we checked that the double depletion was indeed effective in depleting both forms by immunoblotting (Supplementary Figure 1 E).

HP1α accumulates on chromosomes during mitotic exit and forms foci clearly visible in interphase ^33^ (Figure 1 D, yellow arrows). We therefore analysed the number of HP1α foci in the interphase nuclei after H2A.Z.1 or H2A.Z.2 depletion. In this system, we found that only the H2A.Z.2 knockdown, as judged by two independent H2A.Z.2 siRNA oligos, resulted in a significant decrease of the number of HP1α foci (Figure 1 D, E). These differences were not due to a delay of the cell cycle in G1, as cell cycle analyses following H2A.Z.2 knockdown did not show significant changes (Supplementary Figure 1 F). H2A.Z.2 depletion not only did not alter HP1α localisation but also resulted in a different cell cycle profile (Figure 5 D); this aspect will be discussed later in the text.

It is well established that HP1α binds to H3K9me3 to form a repressive chromatin environment ^34, 35^, but recent work indicates that H2A.Z is also able to enhance *in vitro* HP1α binding to nucleosome arrays that do not contain H3K9me3 ^24 25^. If an interplay between H2A.Z, H3K9me3 and HP1 exists *in vivo*, it is not known. However, studies have shown that in mouse cells lacking Ki-67 and presenting a significant decrease in H3K9me3, HP1 maintains its localisation; this suggests that alternative mechanisms exist to maintain HP1-binding at most foci, but the nature of this parallel system is not known. We therefore asked whether H2A.Z could impact on H3K9me3 levels *in vivo*, as this has never been reported. We analysed the levels of H3K9me3 by immunofluorescence after depletion of H2A.Z.1 or H2A.Z.2. Our results showed a significant decrease of H3K9me3 upon H2A.Z.2 siRNA, not only confirming that the variant H2A.Z.2 is involved in heterochromatin regulation, but also suggesting a possible role of H2A.Z.2 upstream or in parallel of H3K9me3 (Figure 1 D, F); however, the enrichment of this marker at the nuclear periphery is still maintained (Supplementary Figure 1 G). We also analysed the levels of another repressive chromatin marker, H3K27me3. H2A.Z.2 depletion increased the levels of H3K27me3 (Supplementary figure 1 H), possibly suggesting that H2A.Z.2 specifically regulates only a subset of heterochromatin. H2A.Z.1 depletion also increased the levels of H3K27me3 with a significantly greater effect than H2A.Z.2 knockdown. This data shows, once again, that H2A.Z paralogues have a different effect on the chromatin landscape.

### H2A.Z.2 is essential for genome stability

A more detailed observation of H2A.Z.2-depleted cells revealed a high incidence of micronuclei, small nuclei formed when a chromosome or a fragment of a chromosome fails to be incorporated into the main cell nuclei after mitosis (Figure 2 A -white arrows). We quantified the number of micronuclei (Figure 2 B) and the number of chromatin bridges and lagging chromatin in anaphase cells (Supplementary figure 2 A, B) upon depletion of each variant: although both H2A.Z.1 and H2A.Z.2 depletion increased the number of micronuclei and anaphase bridges, the phenotype was much stronger in H2A.Z.2-depleted cells. H2A.Z.2 presents two splice variants: H2A.Z.2.1 and H2A.Z.2.2. The latter lacks the utmost C-terminal tail but retains the extended H2A.Z acidic patch. The H2A.Z.2 siRNA oligo used for these experiments targets both H2A.Z.2 isoforms (in the mRNA 5’ UTR), thus we tagged with GFP the coding region of H2A.Z.2.1 and H2A.Z.2.2 cDNA to generate the oligo-resistant mutants (GFP:H2A.Z.2.1^WT^ and GFP:H2A.Z.2.2^WT^) (Figure 2 C). This micronucleation defect was rescued by the GFP:H2A.Z.2.1 construct (but not by the H2A.Z.2.2 one), thus confirming the specificity of the phenotype (Figure 2 D, E). Using a LacO-LacI tethering system that we established in DT40 cells ^36^, we confirmed that the H2A.Z chaperone RFP:LacI:YL1 (part of both the TIP60 and SRCAP complexes) was able to recruit both isoforms to the same extent (Supplementary figure 2 C, D). However, while H2A.Z.2.1 was enriched on mitotic chromosomes, H2A.Z.2.2 was not (Figure 2 D); this was not surprising, since previous studies already reported a partial incorporation of H2A.Z.2.2 into the chromatin ^9^, however our results show that these differences are not due to the ability of binding to the chaperone.

**Figure 2.**
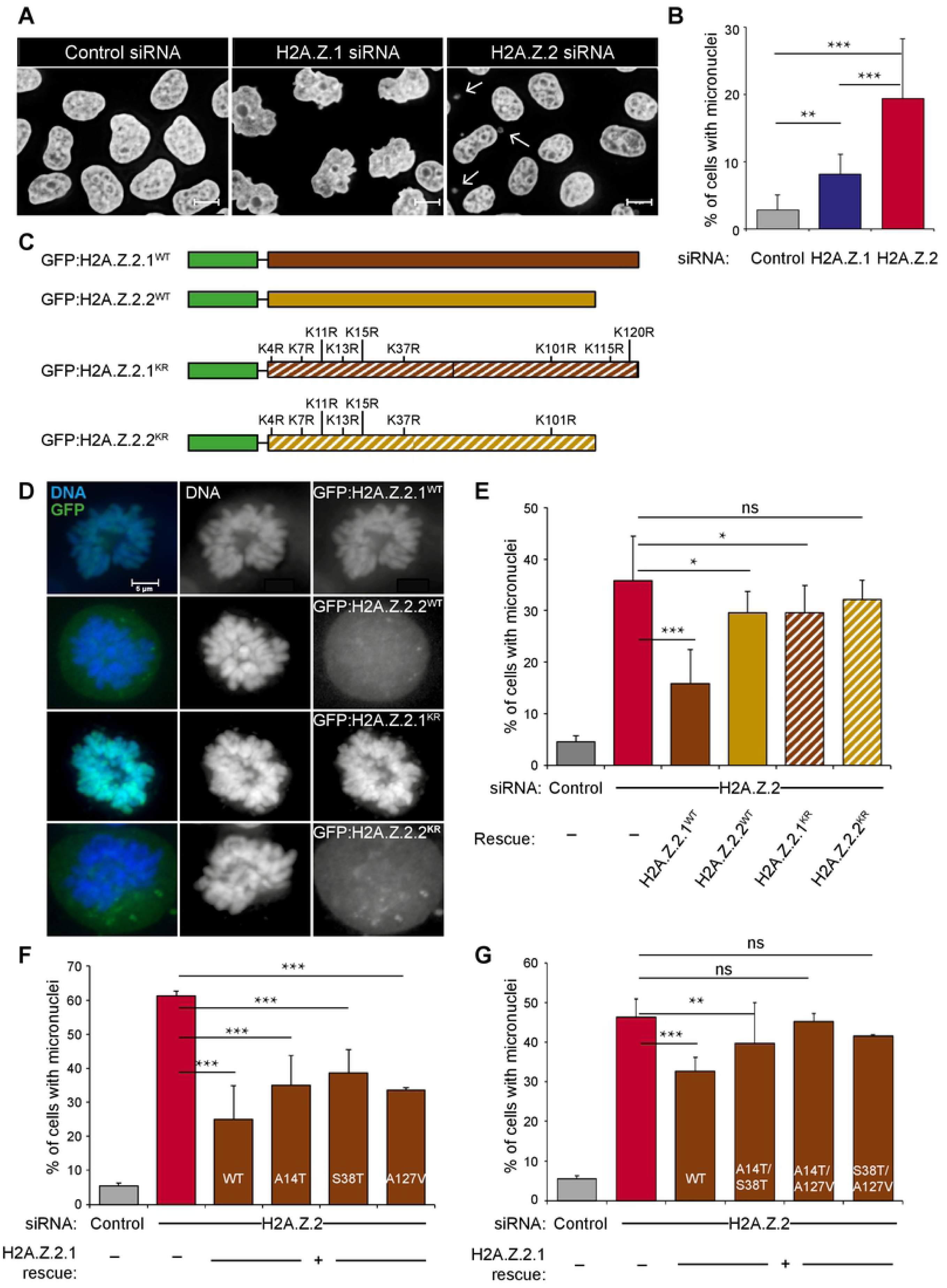
H2A.Z.2.1 knockdown leads to genome instability. **A)** Representative images of HeLa GFP:HP1α cells treated with control, H2A.Z.1 or H2A.Z.2 siRNA for 72 h, fixed, and stained with DAPI. White arrows point at micronuclei. Scale bar: 10 μm. **B)** Quantification of the percentage of cells with micronuclei from experiment in (A). At least 400 cells from three biological replicates were analysed for each condition. The error bars represent the SD. Data sets were statistically analysed using Chi-square test. **=p<0.01; ***=p<0.001. **C**) Schemes of the GFP constructs used for the rescue experiments in (D) and (E). Green boxes represent the GFP, brown boxes represent the H2A.Z.2.1 isoform and yellow boxes represent the H2A.Z.2.2 isoform. Solid fill represents the WT construct whereas striped boxes represent the KR mutant forms. **D)** Representative images of prometaphase chromosomes from HeLa cells co-transfected with H2A.Z.2 siRNA and each of the constructs in C (green). Scale bar: 5μm. **E)** Quantification of the percentage of cells with micronuclei from experiment (D). At least 500 cells were analysed for each category. The error bars represent the SD of three biological replicates. Data sets were statistically analysed using Chi-square test. *=p<0.05; ***=p<0.001; ns= not significant. **F, G)** Quantification of the percentage of cells with micronuclei from HeLa cells co-transfected with H2A.Z.2 siRNA and GFP:H2A.Z.2.1^WT^ with either single (F) or double mutations (G). At least 500 cells were analysed for each category. Error bars represent the SD of three biological replicates. Data sets were statistically analysed using Chi-square test. *=p<0.05; **=p<0.01; ***=p<0.001; ns=not significant. See also Supplementary Figure 2.

As the acetylation status of H2A.Z has been shown to influence its activity in yeast ^37, 38^, we wanted to test the role of acetylation (or other post-translational modifications) for the maintenance of genome stability in mammalian cells. We mutated all the acetylable lysines in both H2A.Z.2.1 and H2A.Z.2.2 to non-acetylable residues (arginines) (GFP:H2A.Z.2.1^KR^ and GFP:H2A.Z.2.2^KR^) (Figure 2 C). The GFP:H2A.Z.2.1^KR^ mutant still localized onto the mitotic chromosomes normally, but GFP:H2A.Z.2.2^KR^ was not, as its wild type counterpart (Figure 2 D). To really demonstrate that GFP:H2A.Z.2.1^KR^ mutant is indeed incorporated into nucleosomes, we prepared mono-nucleosomes from HeLa cells transfected with either GFP:H2A.Z.2.1^WT^ or GFP:H2A.Z.2.1^KR^, separated the histones by SDS PAGE and detected the presence of GFP by immunoblotting (Supplementary figure 2 E, F). These evidence taken together demonstrate that, despite the number of substitutions, the GFP:H2A.Z.2.1^KR^ mutant is indeed incorporated into chromatin.

In order to perform the rescue experiments, we co-transfected H2A.Z.2 siRNA-treated cells with the previously described constructs and quantified the number of micronuclei (Figure 2 E). The data showed that only the GFP:H2A.Z.2.1^WT^ rescues the phenotype in a statistically significant manner, suggesting that H2A.Z.2.1 is important for genome stability maintenance, and that post-translational modifications (acetylation and/or methylation or sumoylation) are also required for this function.

As H2A.Z.1 and H2A.Z.2 only differ by three amino acids, we also asked whether a particular amino acid was crucial for H2A.Z.2 role in genome stability. For this purpose, we mutated the three distinct amino acids of H2A.Z.2.2 back to the one present in H2A.Z.1, either individually or in combination, and performed rescue experiments as described above. From the six mutants, the A14T/A127V and S38T/A127V double mutants failed completely to rescue the micronuclei phenotype (Figure 2 F, G).

In conclusion, our results suggest that the H2A.Z.2.1 variant is playing a key role in chromosome segregation, and that this is also mediated by its post-translational modification status.

### H2A.Z.2 regulates sister chromatids cohesion

We next aimed to understand the molecular mechanisms underlying the micronuclei formation in H2A.Z.2-depleted cells. The presence of micronuclei could be the result of either chromosome mis-segregation or DNA damage arising from double-strand breaks (DSB) and the production of chromosome fragments. To test these hypotheses, we knocked down H2A.Z.2 in a stable cell line that expresses the centromeric histone variant CENP-A tagged with YFP (YFP:CENP-A) and analysed the presence of centromeric signals in the micronuclei. As shown in Figure 3 B, the majority of micronuclei have at least one CENP-A signal, indicating that they possibly contain chromosomes rather than DNA fragments. Mitotic delays have been associated with chromosome segregation errors that might lead to the formation of micronuclei ^39^. For this reason, we investigated the mitotic progression in H2A.Z.2-depleted cells, but we only found a slight decrease in the mitotic index but no significant difference in the distribution of mitotic phases (Supplementary Figure 2 G, H).

**Figure 3.**
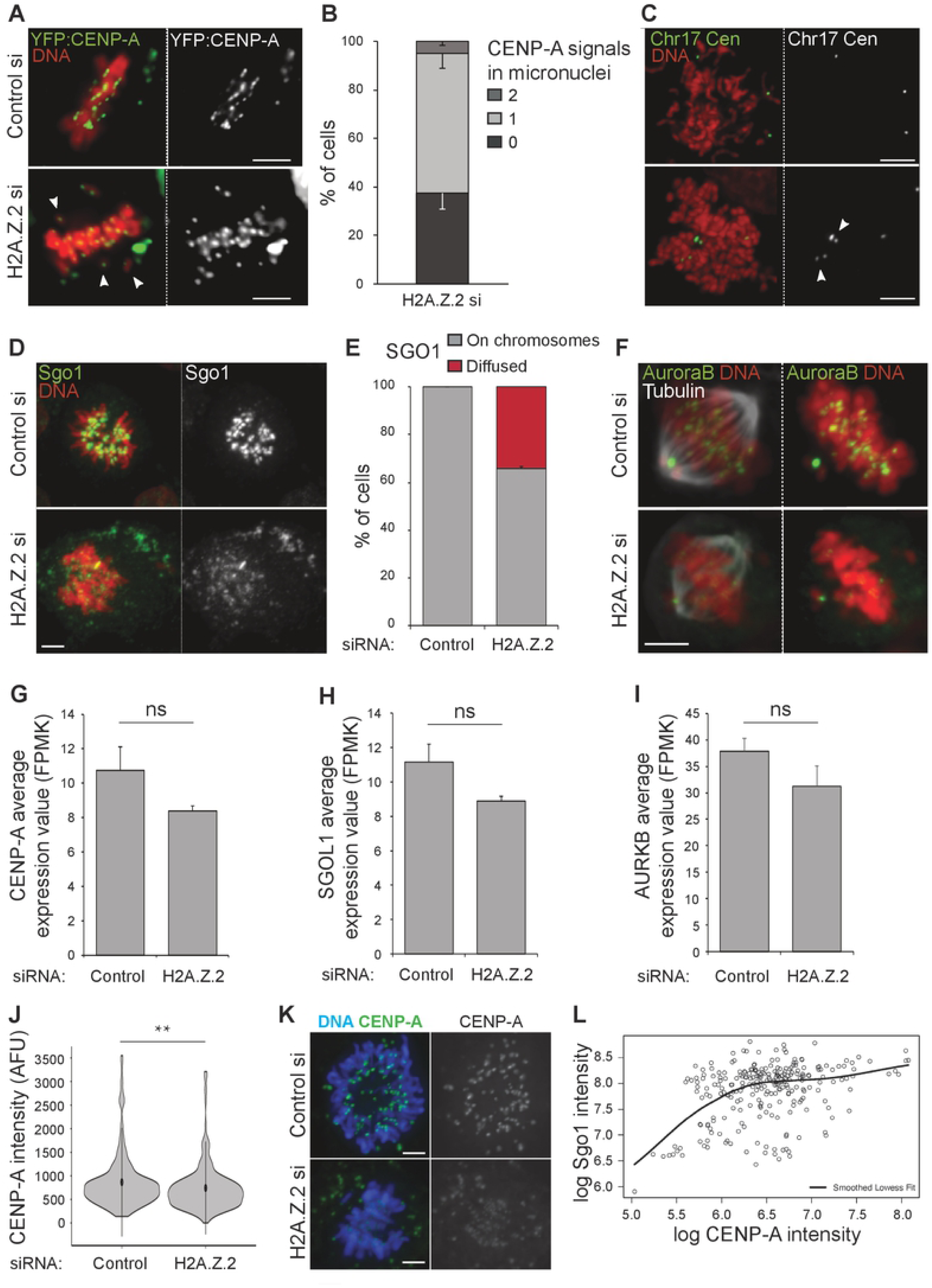
H2A.Z.2 regulates chromosome segregation. **A)** Representative images of metaphases from YFP:CENP-A (green) HeLa cells treated with Control (top) or H2A.Z.2 (bottom) siRNA. The white arrowheads indicate the mis-aligned chromosomes. Scale bar: 10 μm. **B)** Quantification of micronuclei containing 0, 1 or 2 CENP-A signals in HeLa YFP:CENP-A cells transfected with control and H2A.Z.2 siRNA for 72 h. 160 micronuclei were analysed. Error bars indicate SD of three biological replicates. **C)** Representative images of metaphase spreads from control (top) or H2A.Z.2 (bottom) siRNA-transfected HeLa cells after FISH with a Chr17 centromeric probe (green). The arrowheads indicate the separated sister chromatids. Scale bar: 10 μm. **D)** Representative images of HeLa cells transfected with control (top) or H2A.Z.2 (bottom) siRNA, fixed, and stained for Sgo1 (green). Scale bar: 10 μm. **E)** Quantification of Sgo1 localisation in pro-metaphase cells from the experiment in (D). Error bar represents SD of two biological replicates. 35 pro-metaphase cells were analysed. **F)** Representative images of HeLa cells treated as in (D) and stained for Aurora B (green) and alpha tubulin (grey). Scale bar: 10 μm. **G)** *CENP-A* average expression values obtained by RNA sequencing of three biological replicates after control and H2A.Z.2 siRNA treatment. Error bars show the standard deviation (SD). ns=not significant. **H)** *SGOL1* average expression values obtained by RNA sequencing of three biological replicates after control and H2A.Z.2 siRNA treatment. Error bars show the standard deviation (SD). ns=not significant. **I)** *AURKB* average expression values obtained by RNA sequencing of three biological replicates after control and H2A.Z.2 siRNA treatment. Error bars show the standard deviation (SD). ns=not significant. **J)** Violin plot of the CENP-A intensity levels from the centromeres of pro-metaphase and metaphase cells from the experiments in A. 200 cells were analysed. Mean and SD are shown. Data sets were statistically analysed using the Wilcoxon rank test in R. **=p<0.01. **K)** Representative images of HeLa YFP:CENP-A (green) mitotic cells after Control (top) or H2A.Z.2 (bottom) siRNA treatment. Scale bar: 5 μm. **L)** Graph showing the correlation between YFP:CENP-A signals and the intensity of Sgo1 signals at the centromeres of pro-metaphase chromosomes after H2A.Z.2 depletion. The black line represents the lowess smoothed fit.

A compromised Spindle Assembly Checkpoint (SAC) is also compatible with a lack of mitotic arrest and progression through mitosis. Thus, we looked at the CENP-A signals of mis-aligned chromosomes in metaphase: some chromosomes had two CENP-A signals – indicating the presence of a whole chromosome – while others presented a single signal – suggesting that they have been generated by single chromatids (Figure 3 A and B, white arrows). The latter could indicate that the origin of micronuclei in H2A.Z.2-depleted cells is caused by a premature sister chromatid separation, rather than by merotelic attachments or error correction defects. In order to corroborate this observation, chromosome spreads of cells depleted for H2A.Z.2 were subjected to Fluorescence in situ hybridization (FISH) with a probe against the centromeric region of chromosome 17. In control cells, only three FISH signals are observed (the HeLa cell line used has three copies of chromosome 17), but following H2A.Z.2 depletion, the number of FISH signals in mitosis doubled (Figure 3 C) (the modal number of chromosome 17 in interphase is still 3 – Supplementary Figure 3 D, E); this indicates that sister chromatids are prematurely separated in mitosis.

In mitosis, centromeric cohesin removal is prevented until anaphase onset by Shugoshin (Sgo1). In order to investigate whether H2A.Z.2 depletion affected chromosome segregation via altering Sgo1 loading at centromeres, we stained H2A.Z.2-depleted HeLa cells with Sgo1 antibodies. In 40% of pro-metaphase cells, Sgo1 failed to localize to the centromeres and appeared dispersed (Figure 3 D, E). However, the mRNA levels of SGOL1 remained unchanged (Figure 3 H). In a similar way, Aurora B, that together with INCENP, Survivin and Borealin form the chromosomal passenger complex (CPC), was mislocalised in mitosis upon H2A.Z.2 depletion (Figure 3 F) without any significant variation on its mRNA level (Figure 3 I). HP1α, Sgo1 and Haspin are all involved in recruiting the CPC to the centromere ^32, 40-43^. Thus, we asked if depletion of H2A.Z.2 would change the centromeric chromatin and affect centromere maintenance. Analysis of CENP-A levels using the YFP:CENP-A cell line showed that CENP-A is reduced at centromeres of H2A.Z.2-depleted mitotic chromosomes (Figure 3 J, K) but CENP-A mRNA are unchanged (Figure 3 G). Reduced CENP-A levels could be at the basis of a dysfunctional kinetochore and lead to the decrease in cohesin and Aurora B; alternatively, the two pathways could be independent. To better understand this aspect, we analysed the correlation between Sgo1 and CENP-A levels at centromeres of H2A.Z.2-depleted cells: a component on the dependency seems to correlate with CENP-A levels but we cannot exclude that other factors are also playing a role in the amount of Sgo1, particularly at the higher levels (Figure 3 L).

Altogether the data suggest that in human cells the H2A.Z.2.1 paralogue, regulated by its post-translational modification status, is necessary for the maintenance of a functional centromere and its removal affects the recruitment of several chromosome segregation-related proteins to the centromere.

### H2A.Z.1 depletion alters nuclear morphology

H2A.Z.1-depleted cells did not show alterations in HP1α localisation, but they presented an aberrant nuclear shape, a defect not evident in H2A.Z.2-depleted cells (Figure 1 D). To quantify the phenotype, we analysed the nuclear circularity of control, H2A.Z.1- and H2A.Z.2-depleted cells using the NIS Elements AR analysis software (NIKON); the results showed a significant decrease in the circularity of nuclei upon H2A.Z.1 knockdown (Figure 4 A). To confirm that the phenotype was specifically due to H2A.Z.1 depletion and to analyse whether its post-translational modifications were necessary for this function, we generated GFP-tagged oligo resistant construct for H2A.Z.1^WT^ and H2A.Z.1^KR^: both constructs were able to rescue the phenotype (Figure 4 B). These data indicate that H2A.Z.1 is required to maintain a round nuclear shape, but its post-translational modifications are dispensable for this function. To our knowledge, no studies have shown this phenotype in H2A.Z-knockdown cells before. Interestingly, although H2A.Z.2 depleted cells show defects in HP1 localisation, they still maintain a round morphology, almost rounder than control cells (Figure 4 A).

**Figure 4.**
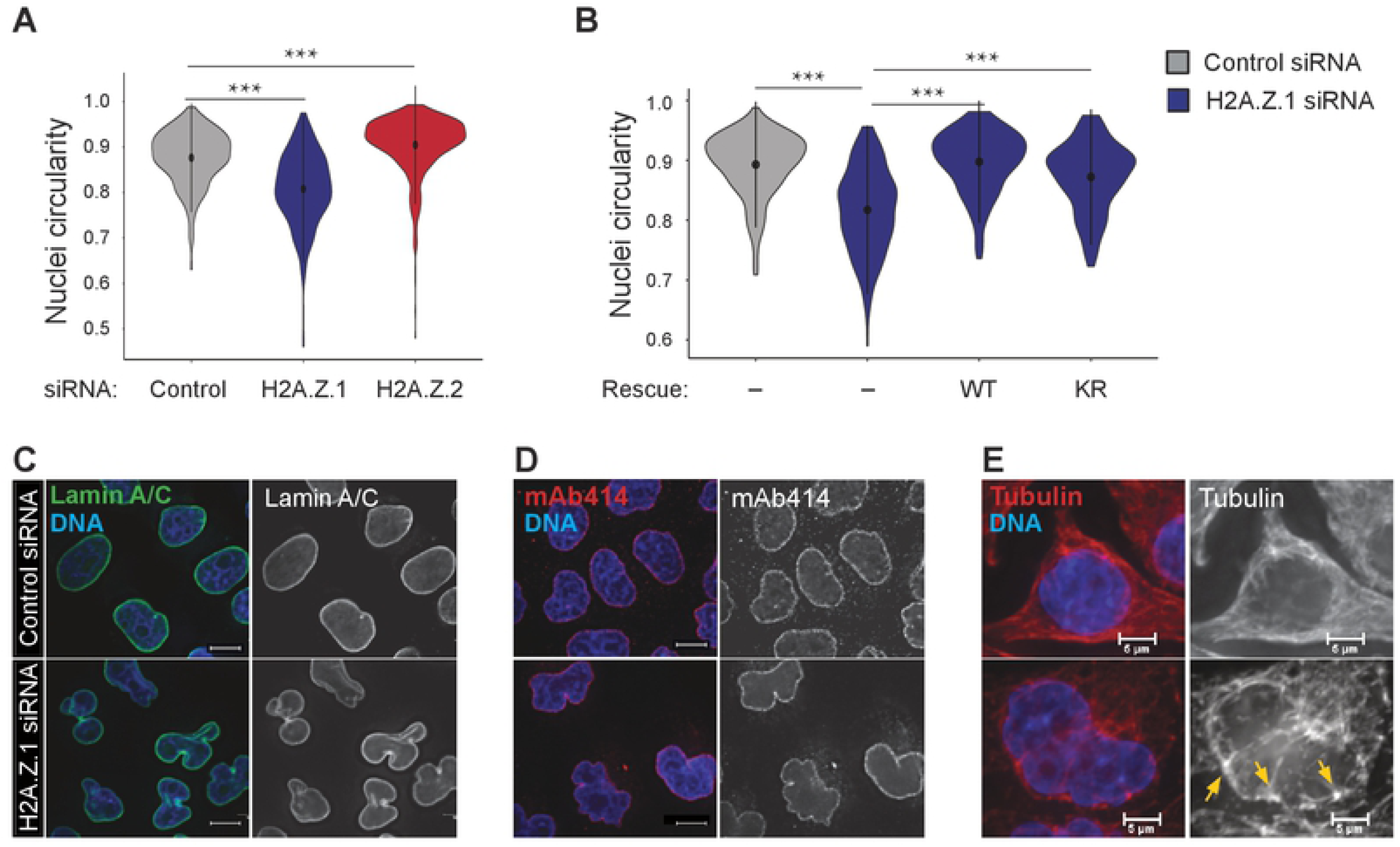
H2A.Z.1 depletion results in aberrant nuclear morphology. **A)** Violin plots of the nuclear circularity index of HeLa cells transfected with control (grey), H2A.Z.1 (blue) or H2A.Z.2 (red) siRNA. The nuclear circularity was analysed using the NIS Elements AR analysis software (NIKON). At least 140 nuclei from three biological replicates were analysed for each condition. Mean and SD are shown. Data sets were statistically analysed using the Wilcoxon rank test in R. ***=p<0.001. **B)** Violin plots of the nuclear circularity index of HeLa cells transfected with control (grey) or H2A.Z.1 (blue) siRNA either alone or in combination with siRNA-resistant GFP:H2A.Z.1^WT^ (WT) or GFP:H2A.Z.1^KR^ (KR) plasmids. The nuclear circularity was analysed using the NIS Elements AR analysis software (NIKON). At least 140 nuclei from three biological replicates were analysed for each condition. Mean and SD are shown. Data sets were statistically analysed using the Wilcoxon rank test in R. ***=p<0.001. **C - E)** HeLa cells transfected with either control (top) or H2A.Z.1 siRNA (bottom) and stained for Lamin A/C (green) (C), mAb4144 (red) for the nuclear pore complexes (D) or alpha-tubulin (red) (E). Yellow arrows (E) point at accumulation of tubulin at invaginations. Scale bar: 10μm (C, D) and 5μm (E).

**Figure 5.**
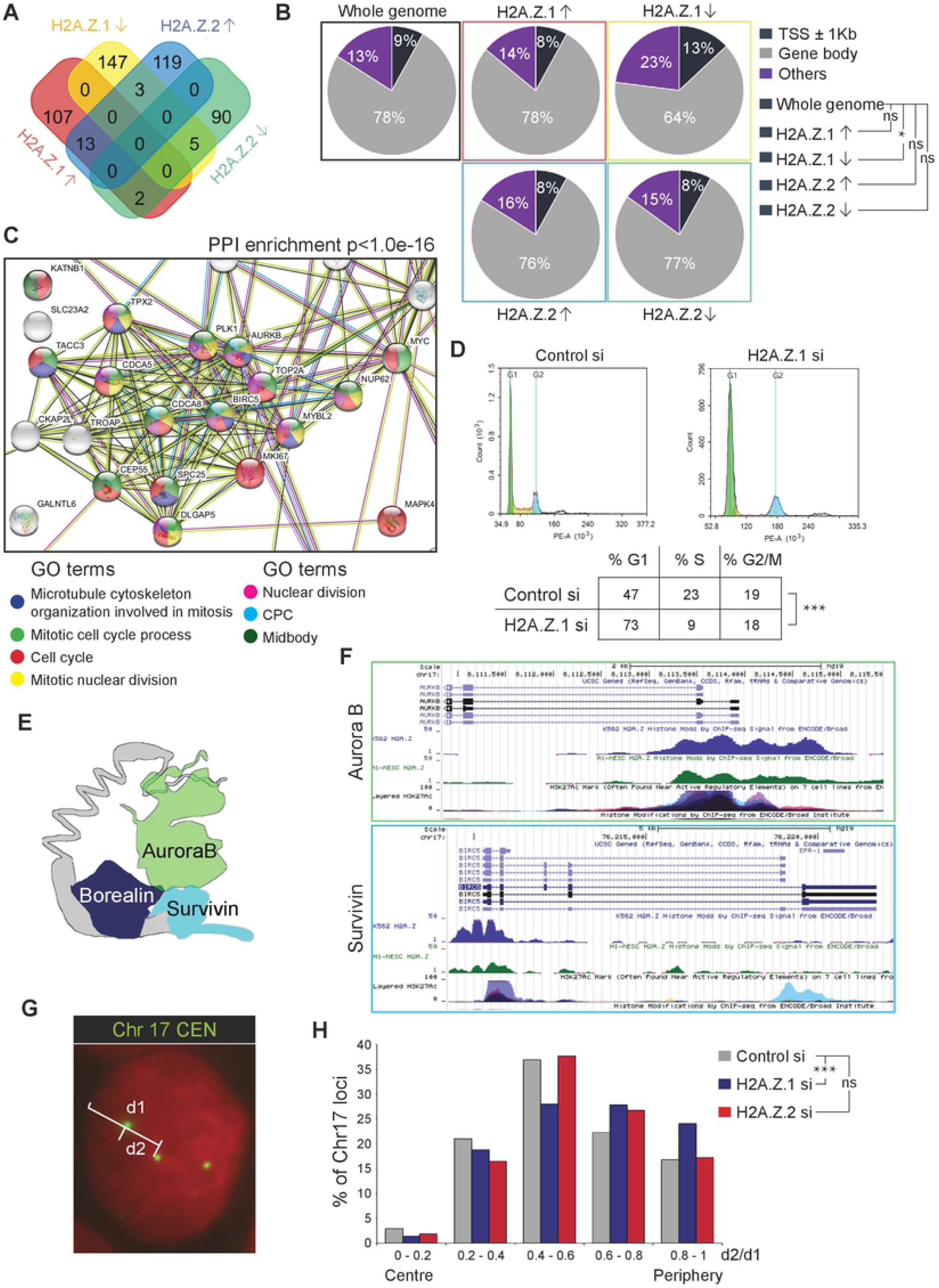
H2A.Z.1 regulates the expression of cell cycle-related genes. **A)** Gene expression of three biological replicates of HeLa cells transfected with control, H2A.Z.1 (blue) or H2A.Z.2 siRNA was analysed by RNA sequencing. Venn diagram shows the number of significant (p<0.01) differentially expressed genes and the overlap between each set of genes. The upwards arrows indicate up-regulation and the downward arrows indicate down-regulation. **B)** Pie chart displays the percentages of H2A.Z peaks at or nearby transcription start sites (TSS) (dark blue), within the gene body (grey) or elsewhere in the whole genome (purple) for the differentially expressed genes following H2A.Z.1 or H2A.Z.2 depletion. The upwards arrows indicate up-regulation and the downward arrows indicate down-regulation. A 2-sample test for equality of proportions was used for the statistical analyses. **=p<0.01; *=p<0.05; ns=not significant. **C)** Downregulated genes following H2A.Z.1 depletion were analysed by STRING. The image shows the cluster of genes with gene onthology (GO) terms related to cell cycle. PPI=Protein-protein interaction. **D)** Flow cytometry analyses profiles of control and H2A.Z.1 siRNA-treated HeLa cells. Percentages represent the mean of two biological replicates. Data sets were statistically analysed using Chi-square test. ***=p<0.001. **E)** Cartoon showing the altered members of the Chromosome Passenger Complex (CPC) after H2A.Z.1 depletion. **F)** UCSC analyses of H2A.Z localisation on Aurora B (green) and Survivin (blue) genes showing H2A.Z enrichment at the TSS. **G)** Representative image of a HeLa nucleus after FISH with Chr17 centromeric probe (green). The distance of the FISH signals from the periphery was calculated as follows: the distance from the centre of the nucleus to the periphery (d1) and the distance from the centre of the nucleus to the FISH signal (d2) were measured. **H)** Distribution of the position within the nucleus of the centromere of Chr17 from the experiment in (G) calculated as the ratio d2/d1. At least 500 nuclei were analysed per condition. Data sets were statistically analysed using Chi-square test. ***=p<0.001. See also supplementary figure 3.

Alterations in chromatin organisation have been associated with nuclear rigidity and nuclear blebs formation ^44^. For example, alterations in histone modifications resulted in nuclear blebs that were depleted for heterochromatin but enriched for euchromatin ^45^. Although we did not see a decrease in global H3K9me3 upon H2A.Z.1 depletion (Figure 1 F), the phenotype on nuclear morphology could be explained if the normal enrichment of this histone mark at the nuclear periphery was compromised. However, H3K9me3 enrichment at the nuclear periphery remained unchanged (Supplementary Figure 1 G).

Perturbations of the lamina, including lamin A and B1 depletion and lamin A mutations, have also been linked to alterations in nuclear morphology ^46-48^. Thus, we analysed by immunofluorescence both the nuclear lamina and the nuclear pore complexes (NPCs) with anti-lamin A/C and mAb414 (antibody that recognizes a set of FG repeat-containing nucleoporins) antibodies. Lamin A/C and the NPC remained unchanged, at least at this level of resolution, after H2A.Z.1 depletion (Figure 4 C, D).

Microtubules are yet other major effectors impacting on nuclear morphology with a number of studies reporting their importance for normal nuclear shape in interphase ^49, 50^. We analysed α-tubulin distribution in the H2A.Z.1-depleted cells and observed an accumulation of tubulin at the points of invaginations (Figure 4 E, yellow arrows). Although more investigations will be required to unveil the underlying mechanisms, this data suggests that unbalanced mechanical forces possibly account for the nuclear deformity in the H2A.Z.1-knockdown cells.

### H2A.Z.1 and H2A.Z.2 have distinct roles in gene expression regulation

Several studies have linked H2A.Z to transcription regulation ^13-17, 28^; whether it promotes or represses gene expression still remains controversial. These differences seem to depend on the targeted gene and post-translational modifications of H2A.Z itself. A recent study ^30^ analysed the differences between H2A.Z.1 and H2A.Z.2 on gene regulation in U2OS and WI38 human primary fibroblast cells. The data showed that H2A.Z.1 and H2A.Z.2 regulate different sets of genes in different cell types and that, depending on the gene, H2A.Z.1 and H2A.Z.2 either had similar or antagonistic functions ^30^. In order to investigate the differences in transcription following H2A.Z.1 or H2A.Z.2 depletion in our system, we conducted RNA-seq analyses in HeLa cells. Our results revealed a striking difference in gene expression changes between the two variants. In order to narrow down the number of targets, we focused only on genes showing a high statistical significant change in expression with p < 0.01 and fold change > 2. H2A.Z.1 depletion resulted in 147 down-regulated genes and 107 up-regulated genes; H2A.Z.2 depletion resulted in 90 down-regulated genes and 119 up-regulated genes. Among the down-regulated genes, only 5 were common between variants, while among the up-regulates genes, only 13 (Figure 5 A, Supplementary figure 3 A). Our results therefore suggest non-overlapping roles for these two paralogues also in gene regulation.

Based on published ChIP-seq datasets (no antibody is able to discriminate between the two isoforms and therefore ChIP-seq datasets contain genomic regions occupied by either isoform), we compared the distribution of H2A.Z at three different regions (i-the gene body, ii-at transcription start sites ± 1Kb (TSS) and iii-other parts of the gene) for the whole genome and the four different groups of genes that change expression upon H2A.Z.1 or H2A.Z.2 depletion: upregulated or downregulated after H2A.Z.1 depletion and upregulated or downregulated after H2A.Z.2 depletion. Interestingly, H2A.Z is enriched at the TSS of genes that are downregulated only after H2A.Z.1 depletion (Figure 5 B). This possibly suggests that, in cycling cells, H2A.Z.1 has a more pronounced role in maintaining an open chromatin at active promoters than H2A.Z.2.

Using STRING analyses, we identified a highly significant protein-protein interaction (PPI) enrichment (p = 10^−16^) in the downregulated genes of H2A.Z.1-depleted cells – this is the only group that showed a significant PPI enrichment. Most of these genes are linked to gene ontologies (GO) terms associated with cell division and mitosis, and include MYC, MKI67 and AURKB (Figure 5 C). This is in agreement with the flow cytometry analyses of the cell cycle, which revealed a significant increase of cells in G1 phase upon H2A.Z.1 depletion (Figure 5 D), and with a decreased mitotic index in H2A.Z.1-depleted cells (Supplementary Figure 2 G). Three out of the four Chromosome Passenger Complex (CPC) components were significantly downregulated following H2A.Z.1 knockdown (Aurora B, Borealin and Survivin). Furthermore, analysis of ENCODE/Broad Chip-Seq data as reported in UCSC genome browser shows that H2A.Z is indeed mainly localized at their promoters (Figure 5 E, F, Supplementary Figure 3 B). Similarly, CDCA2 (also known as Repo-Man), which counteracts the Aurora B in mitosis and is co-regulated with it via FOXM1 ^51^, is also downregulated upon H2A.Z.1 depletion and, similarly to Aurora B, its TSS is enriched for H2A.Z (Supplementary Figure 3 C). These results indicate that H2A.Z.1 in HeLa cells is involved in cell cycle progression by affecting the expression of several important cell cycle-related genes. This phenotype is quite distinct from the one obtained upon H2A.Z.2 depletion where neither changes in cell cycle profile (Supplementary Figure 1 F) nor significant decrease in Aurora B mRNA (Figure 3 I) were observed.

Several genes differentially expressed upon H2A.Z.1 knockdown are localised on chromosomes 1 (p<0.05), 17 (p<0.05), and 22 (p<0.05) (Supplementary figure 3 F) while only a single significant enrichment on chromosomes 9 (p<0.05) is obtained for genes affected by H2A.Z.2 knock-down. Chromosomes occupy specific territories in the nucleus that are characteristic for each cell type and linked to gene expression ^52, 53^. In order to assess if chromosome positioning could have been affected by H2A.Z.1 depletion, we analysed the localisation of chromosome 17 using a centromeric FISH probe (Figure 5 G, H and supplementary Figure 3 D): high d2/d1 ratios represent signals located at the nuclear periphery, while low d2/d1 ratios represent signals located towards the centre of the nucleus. The results show that following H2A.Z.1 depletion, chromosome 17 is moved towards the nuclear periphery and could explain the repression of many genes located on this chromosome (Figure 5 H). This change in position does not occur upon depletion of H2A.Z.2 (Figure 5 H).

We suggest that H2A.Z.1 and H2A.Z.2 regulate the expression of different subsets of genes and that H2A.Z.1 is particularly involved in cell cycle progression possibly via the maintenance of an open chromatin at the promoters of key cell-cycle genes.

## DISCUSSION

Despite the established importance of the histone variant H2A.Z in many aspects of cell biology and growing evidence suggesting differential roles for the H2A.Z.1 and H2A.Z.2 paralogues in chromatin organisation and function, the majority of the studies thus far have focussed on H2A.Z as a single variant and very little is known about their individual roles (if any) during the cell cycle in mammals. In this study we have shown specific and distinct functions of H2A.Z.1 and H2A.Z.2 in heterochromatin regulation, chromosome segregation and cell cycle progression in human cells.

### H2A.Z.2 is essential for chromosome segregation

Using an unbiased proteogenomic approach, we have shown that HP1α preferentially binds to H2A.Z-enriched chromatin *in vitro* and that specifically the histone variant H2A.Z.2 is necessary for heterochromatin maintenance in human cells. Although some links between H2A.Z and heterochromatin regulation had been suggested ^24, 25, 54-56^, a specific role for one of the two H2A.Z paralogues had never been reported. Here we show that only H2A.Z.2 depletion decreases the number of HP1α foci and H3K9me3 levels in interphase in human cells. H2A.Z.2-specific depletion in HeLa cells resulted in profound chromosome segregation errors that led to micronuclei formation. Our study has provided clear evidence for the different roles of the splice variants and their post-translational modifications that are essential for chromosome segregation. The isoform H2A.Z.2.1, but not its shorter variant H2A.Z.2.2, is critical for chromosome segregation and this function depends on its post-translational modification status, since we have demonstrated that both the wild-type and mutant forms can be incorporated into the chromatin with the same efficiency, but the mutant cannot rescue the segregation defect phenotype. Although the shorter H2A.Z.2.2 variant possesses the acidic patch responsible for its deposition, it lacks the C-terminal tail, which has been shown to be important in Drosophila ^57^ and yeasts ^58^. Although the specific mechanism is not known, the C-terminal tail contains several residues that can be post-translational modified and could potentially have an impact on the stability of the nucleosomes.

As H2A.Z.1 and H2A.Z.2 differ by only three amino acids, we have also investigated which one was critical for the mitotic function. Single amino acid mutations – substituting one of the three different amino acids of H2A.Z.2 to H2A.Z.1 – were all able to rescue the micronuclei phenotype, but a double mutation of Alanine 14 to Threonine and Threonine 38 to Serine was not. This indicates that both amino acids 14 and 38 confer H2A.Z.2 with the ability to regulate chromosome segregation. Amino acid 38 is important for the L1 loop structure and nucleosome stability ^8^ and the S38T substitution in H2A.Z.1, mimicking T38 found in H2A.Z.2, can rescue the SRCAP-dependent Floating-Harbor Syndrome ^27^ defects. The role of the amino acid 14 is still unknown but being an alanine that cannot be phosphorylated, (compared to the threonine in the paralogue counterpart), it could suggest that a differential phosphorylation between the two forms may play a critical role, especially during mitosis. In our experiments all these mutants were efficiently incorporated into the chromosomes, suggesting that the lack of rescue can be linked to either the ability of generating a different chromatin conformation or to the recruitment of specific effectors.

We have also investigated the molecular mechanisms behind the chromosome segregation defects. H2A.Z.2-depleted cells presented impaired loading of CENP-A, Sgo1 (protector of centromeric cohesion) and Aurora B (member of the CPC). Interestingly, two H2A.Z specific peptides (GDEELDSLIK and ATIAGGGVIPH – note that these sequences are present in both H2A.Z.1 and H2A.Z.2) were enriched in the CENP-A nucleosome pull-downs when compared with those containing histone H3.1 ^59^. This suggests that a specific centromeric CENP-A containing chromatin could favour the maintenance or deposition of CENP-A. Interestingly, the majority of H2A.Z deposition at pericentric heterochromatin occurs in G1 ^60^, at the same time as CENP-A. Treatment with 5’-aza 2’-deoxycytidine (5-Aza) leads to an increased incorporation of both CENP-A and H2A.Z at pericentric heterochromatin that has been interpreted as a consequence of a disruption in the heterochromatin at the centromere ^60^. However, our data seems to suggest that H2A.Z.2 incorporation is upstream and can affect CENP-A deposition or stability. This is a novel and interesting aspect that will be further investigated.

We have also observed a decrease in Aurora B levels at centromeres of mitotic chromosomes in cells depleted of H2A.Z.2. It was previously reported that INCENP^360-876^ could interact *in vitro* with H2A.Z and that the GDEELDSLIKA region was essential for the binding ^55^. This could explain the results we have obtained, but it does not account for the specificity; in fact, the interacting region is the same in both paralogues.

Our results have also shown a link between H2A.Z.2 and sister chromatid cohesion. This effect could be possibly mediated by the impairment of HP1α localisation. HP1α is removed from the chromatin during mitosis, but a fraction remains at pericentric heterochromatin where it is involved in the recruitment of cohesin and Sgo1 ^41, 59, 61-64^. However, HP1 also plays important roles in the regulation of components that support kinetochore function (the CPC) prior to and during mitotic entry, where HP1-induced CPC clustering appears to be an effective way of promoting the activation of Aurora B kinase ^43^.

Finally, the data we have provided on the correlation between CENP-A and the Sgo1 levels at centromere may suggest that H2A.Z.2 is a key chromatin player at the intersect of different pathways that altogether contribute to maintain a functional centromere.

### H2A.Z.1 is important for the maintenance of nuclear morphology

One of the most surprising observations upon H2A.Z.1 depletion was the abnormal nuclear morphology. Perturbations in the lamina and chromatin organization have been linked to nuclear morphology alterations ^45, 65-67^, but both the lamina and heterochromatin organization remained unchanged following H2A.Z.1 depletion in our system.

Cytoskeleton forces have also been associated with nuclear morphology maintenance ^47, 68-71^ and studies in Drosophila have shown that the H2A.Z orthologue, H2A.V, promotes microtubule formation ^72^. Microscopic observations showed accumulation of α-tubulin in the nuclear invaginations of H2A.Z.1-depleted cells, although further studies would be needed to understand the underlying mechanisms. Experiments in *Xenopus* eggs have shown that inhibition of microtubule polymerisation by Dppa2 is necessary for normal nuclear shape ^49^, but analyses of our RNA-seq dataset did not show a significant change on *DPPA2* expression upon H2A.Z.1 depletion. Although these observations are still descriptive at the present stage, they indicate a distinct role for H2A.Z.1 in genome organisation. Future studies aimed at understanding the overall chromatin structure caused by H2A.Z.1 depletion and specific H2A.Z.1 interactors could reveal the mechanisms at the basis of these abnormalities.

### H2A.Z.1 and H2A.Z.2 have separate functions on gene expression regulation

Our RNA-seq analyses also revealed a completely different role for H2A.Z.1 and H2A.Z.2 in transcription regulation. These distinct effects could be due to the different binding partners reported for H2A.Z.1 and H2A.Z.2: H2A.Z.1 has preference for interacting with BRD2, PHF4, HMG20A and TCF20, whereas H2A.Z.2 has been shown to interact with SIRT1 ^73 74 30^. A recent study conducted in U2OS and fibroblast cells analysed the target transcriptomic landscape after individual depletion of each H2A.Z variants; the results showed both distinct and overlapping sets of genes, as well as similar or antagonistic functions, depending on the targets ^30^. Our datasets differ from the reported one as we obtained an almost non-overlapping picture of gene expression changes and we observed a significant enrichment of H2A.Z at the TSS of genes that undergo downregulation after H2A.Z.1 depletion. In agreement with this study, our data has shown that H2A.Z.1 depletion resulted in downregulation of a subset of cell-cycle regulated genes, which explains the cell cycle arrest and low cell division rate we have reported here. However, Lamaa and colleagues ^30^ also found downregulation of cell cycle-related genes after H2A.Z.2 depletion; we did not see this in our system.

Our findings show that H2A.Z.1 is crucial for cell cycle progression via the regulation of many cell cycle-related genes. This is in agreement with the observed overexpression of H2A.Z in a variety of malignant tumours, including breast ^75^, prostate ^76^ and bladder ^77^ cancers and metastatic melanoma ^28^. In particular, H2A.Z.1 has been linked to liver tumorigenesis where its depletion induced the expression of several cell cycle inhibitors ^78^. Moreover, the co-regulation of the CPC and its counteracting phosphatase Repo-Man (CDCA2) by H2A.Z.2 we discovered, provides further support to the importance of the role of these complexes in cancer where the co–upregulation of Repo-Man and Aurora B in tumours is inversely correlated with patient survival ^51^. This again underlines the potential importance of H2A.Z.1 for cancer progression.

Interestingly, a few studies have pointed at the possibility that it is the balance between the two paralogues that allows for a normal cell proliferation. In this respect, as we have shown that H2A.Z.2 depletion resulted in chromosome instability while H2A.Z.1 supports the expression of master cell cycle genes, we can envisage that slight changes in their balance could easily produce an aberrant cancer prone phenotype, on one hand by supporting proliferation and on the other by increasing genome instability, both beneficial for a tumour cell.

In conclusion, we have demonstrated non-redundant roles for H2A.Z.1 and H2A.Z.2 in different aspects of cell biology and gene expression, highlighting the importance of studying these variants independently. More studies will require detailed analyses of the downstream effectors of these paralogues, the chaperones and machineries dedicated to the specific deposition of these variants. This is particularly important for therapeutic purposes, as only targeting the correct variant would offer a proper intervention.

## MATERIALS AND METHODS

### Cell culture, cloning and transfections

HeLa cells were grown in DMEM supplemented with 10% foetal bovine serum (FBS) and 1% Penicillin–Streptomycin (Invitrogen Gibco) at 37°C with 5% CO_2_.

DT40 cells carrying a single integration of the LacO array ^79^ were cultured in RPMI1640 supplemented with 10% FBS, 1% chicken serum and 1% Penicillin–Streptomycin at 39°C and 5% CO_2_.

The HeLa GFP:HP1α cell line was generated by transfection of the GFP:HP1α construct (kind gift from Schirmer lab, Wellcome Trust Edinburgh) with Polyplus JetPrime® reagent transfection and selection in G418 (2 mg/ml).

The HeLa CENP-A:YFP cell line was kindly provided by Dr Lars E.T. Jansen (Oxford University, UK).

Transient transfections for DT40 in LacO array background were conducted as previously described using GFP-fused H2A.Z.1, H2A.Z.2.1 and H2A.Z.2.2 ^79^.

For siRNA treatments, HeLa cells were seeded in 6-well plates, transfected using Polyplus JetPrime® (PEQLAB, Southampton, UK) with the appropriate siRNA oligonucleotides (50 nM), and analysed after 72 h. The siRNAs were obtained from Merck. Control: 5’-CGUACGCGGAAUACUUCGA-3’; H2A.Z.1: 5’-GCCGUAUUCAUCGACACCU-3’; H2A.Z.2: 5’-AUUUGUAUGUUCUUAGACU-3’; H2A.Z.2_2: 5’-GUGACAGUUGUGUGUUGAU-3’.

For rescue experiments, 1 µg of GFP:H2A.Z.1 DNA or 100 ng of GFP:H2A.Z.2 DNA were used. GFP:H2A.Z.1^WT^, GFP:H2A.Z.2.1^WT^, GFP:H2A.Z.2.2^WT^, GFP:H2A.Z.1^KR^, GFP:H2A.Z.2.1^KR^ and GFP:H2A.Z.2.2^KR^ were synthetized by ProteoGenix (La Haye, France). They were all cloned into pEGFP-C1 by XhoI/ KpnI, except GFP:H2A.Z.2.1^WT^ and GFP:H2A.Z.2.2^WT^ that were cloned by BglII/ BamHI1 and BglII/ EcoRI respectively. Single or double mutations were produced with the Q5® Site-directed Mutagenesis kit (New England Biolabs, UK) using the primers in Table 1.

**Table 1.**
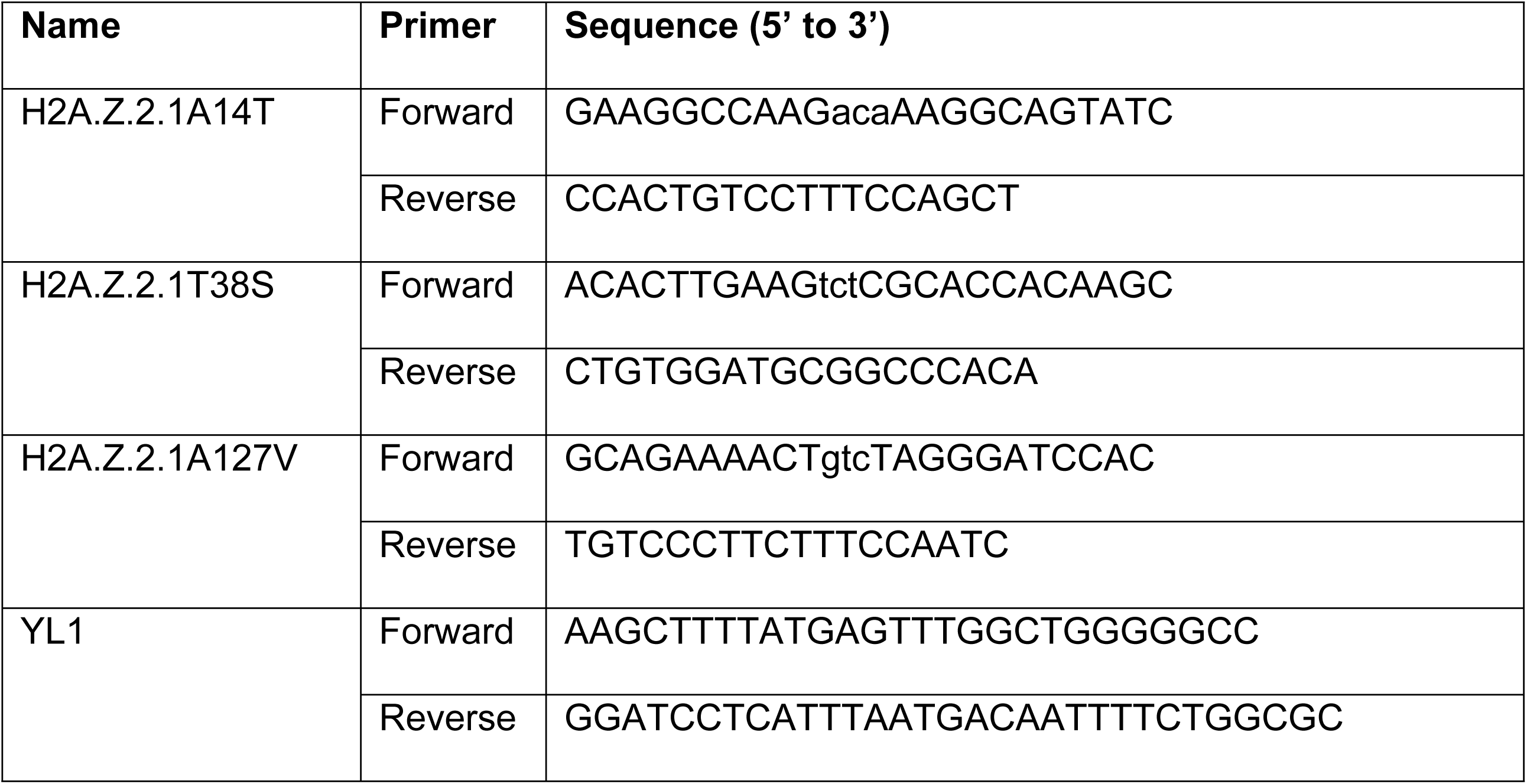
Primers used for cloning

The oligo-resistant H2A.Z.1 mutants were generating by mutating the following oligo target sequence GGC(G) CGT(R) ATT(I) CAT(H) CGA(R) CAC(H) CTA(L) to GGC(G) <u>A</u>G<u>G</u>(R) AT<u>C</u>(I) CAT(H) <u>A</u>G<u>G</u>(R) CAC(H) CTA(L).

To produce the RFP:LacI:YL1 construct, we first generated a GFP:LacI:YL1 plasmid. The YL1 sequence was obtained by PCR using the primers in Table 1 and cloned into GFP:LacI ^31^ by HindIII/BamHI. GFP was replaced by RFP using NheI/BglII.

The primers used for the study were acquired from Eurofins Genomics (Germany) and all the restriction enzymes from New England Biolabs (UK).

### Immunofluorescence microscopy

Cells were fixed in 4% PFA and processed as previously described (Vagnarelli et al., 2006). Primary and secondary antibodies were used as in Table 2. Fluorescence-labelled secondary antibodies were applied at 1:200 (Jackson ImmunoResearch). Three-dimensional data sets were acquired using a wide-field microscope (NIKON Ti-E super research Live Cell imaging system) with a numerical aperture (NA) 1.45 Plan Apochromat lens. The data sets were deconvolved with NIS Elements AR analysis software (NIKON). Three-dimensional data sets were converted to Maximum Projection using the NIS software, exported as TIFF files, and imported into Adobe or Inkscape for final presentation.

**Table 2.** Antibodies used for immunofluorescence

| **Antibody** | **Source** | **Cat number** | **Dilution** |
| --- | --- | --- | --- |
| H3K27me2/3 | Active Motif | #39535 | 1:500 |
| H3K9me3 | Active Motif | #39161 | 1:500 |
| Lamin A+Lamin C | Abcam | #ab108595 | 1:2500 |
| mAb414 | Covance | #MMS-120R | 1:500 |
| Alpha-Tubulin | Sigma | #T5168 | 1:1000 |
| Shugoshin | Abcam | #ab58023 | 1:200 |
| Aurora B/ AIM1 | Cell signalling<br>Technology | #3094 | 1:200 |

For the analyses of the mitotic index, cells stained with DAPI and alpha tubulin were used. At least 500 cells per experimental condition for each biological replica were analysed. Chromosome condensation and spindle morphology were used to categorise the different mitotic stages.

For quantification of the staining, masks were created around the DAPI nuclei using NIS Elements AR analysis software. Mean intensity of antibodies signals were extracted, exported to Excel (MS office) and background was subtracted.

For quantification of enrichment at the LacO locus, four circles were designed: one around the LacI spot, two within the nucleus and one outside the cell. Signals intensities were extracted and the mean of the two circles within the nucleus calculated. The outside circle was used as background and subtracted from both the mean nuclear and the LacI spot, then the signal intensity from the LacI was normalized relative to the intensity of the nuclear signal.

Violin plots were generated using the ggplot2 package in R.

### Preparation of HP1α-bound nucleosomes and mass spectrometry

Preparation of HP1α-bound nucleosomes and mass spectrometry were performed as previously described in de Castro et al., (2017) with the following modifications. Two HeLa cultures were grown in SILAC in RPMI(-)Arg(-)Lys supplemented with 10% dialyzed FBS and either with 100 ng/ml of 13C Lysine plus 30 ng/ml of 13C Arginine (Cambridge isotope Laboratories) for heavy labelling, or 12C Lysine plus 12C Arginine for light labelling. Chromatin was pre-cleared on GST beads and separated on SDS PAGE together with the elution samples from the binding experiments and used as “input control”.

Excised gel bands were stained with Imperial Stain (Thermo Fisher Scientific, UK) de-stained overnight and proteins were digested with trypsin, as previously described ^80^. In brief, proteins were reduced in 10 mM dithiothreitol (Sigma Aldrich, UK) for 30 min at 37°C and alkylated in 55mM iodoacetamide (Sigma Aldrich, UK) for 20 min at ambient temperature in the dark. They were then digested overnight at 37°C with 13 ng μL-1 trypsin (Pierce, UK).

Following digestion, SILAC-labelled samples were mixed with equal amount of the input sample and they were diluted with equal volume of 0.1% Trifluoroacetic acid (TFA) (Sigma Aldrich, UK) and spun onto StageTips as described by Rappsilber et al ^81^. Peptides were eluted in 40 μL of 80% acetonitrile in 0.1% TFA and concentrated down to 5μL by vacuum centrifugation (Concentrator 5301, Eppendorf, UK). The peptide sample was then prepared for LC-MS/MS analysis by diluting it to 5 μL by 0.1% TFA. MS-analyses were performed on a Q Exactive mass spectrometer (Thermo Fisher Scientific, UK) and on an Orbitrap FusionTM LumosTM TribridTM mass spectrometer (Thermo Fisher Scientific, UK), both coupled on-line, to Ultimate 3000 RSLCnano Systems (Dionex, Thermo Fisher Scientific). In both cases, peptides were separated on a 50 cm EASY-Spray column (Thermo Fisher Scientific, UK) assembled in an EASY-Spray source (Thermo Fisher Scientific, UK) and operated at a constant temperature of 50 °C. Mobile phase A consisted of water and 0.1% formic acid (Sigma Aldrich, UK); mobile phase B consisted of 80% acetonitrile and 0.1% formic acid. The total run time per sample was 220 min. In both mass spectrometers, the peptides were loaded onto the column at a flow rate of 0.3 μL min-1 and eluted at a flow rate of 0.2 μL min-1 according to the following gradient: 2 to 40% buffer B in 180 min, then to 95% in 16 min. For the Q Exactive, FTMS spectra were recorded at 70,000 resolution (scan range between 350 and 1400 m/z) and the top 10 most abundant peaks with charge ≥ 2 and isolation window of 2.0 Thomson were selected and fragmented by higher-energy collisional dissociation ^82^ with normalised collision energy of 25. The maximum ion injection time for the MS and MS2 scans was set to 20 and 60 ms respectively and the AGC target was set to 1 E6 for the MS scan and to 5 E4 for the MS2 scan. Dynamic exclusion was set to 60 s. For the Orbitrap FusionTM LumosTM, survey scans were performed at resolution of 120,000 (scan range 350-1500 m/z) with an ion target of 4.0e5. MS2 was performed in the Ion Trap at a rapid scan mode with ion target of 2.0E4 and HCD fragmentation with normalized collision energy of 27 ^82^. The isolation window in the quadrupole was set at 1.4 Thomson. Only ions with charge between 2 and 7 were selected for MS2.

The MaxQuant software platform ^83^ version 1.5.2.8 was used to process the raw files and search was conducted against Homo sapiens complete/reference proteome set of UniProt database (released on 16/03/2017), using the Andromeda search engine ^84^. For the first search, peptide tolerance was set to 20 ppm while for the main search peptide tolerance was set to 4.5 pm. Isotope mass tolerance was 2 ppm and maximum charge to 7. Digestion mode was set to specific with trypsin allowing maximum of two missed cleavages. Carbamidomethylation of cysteine was set as fixed modification. Oxidation of methionine and acetylation of the N-terminal were set as variable modifications. Multiplicity was set to 2 and for heavy labels Arginine 10 and Lysine 8 were selected. Peptide and protein identifications were filtered to 1% FDR. The Perseus computational platform ^85^ version 1.6.0.2 was used for the statistical analysis of the MaxQuant-generated datasets and for the creation of the volcano plots.

The mass spectrometry proteomics data have been deposited to the ProteomeXchange Consortium via the PRIDE partner repository with the dataset identifier PXD018719.

### FISH

FISH was performed in HeLa cells as previously described ^31^ using a probe against chromosome 17 ^86^.

For the centromere position analyses, a single plane containing each spot was selected. The distance of the spots from the periphery was measured and represented as a fraction of the radius on which each spot belongs.

### Cell cycle analysis with flow cytometry

Cells were trypsinised, resuspended and incubated at room temperature for 30 min in 70% ice-cold ethanol. Cells were centrifuged at 1000 g for 4 min, washed with PBS and the supernatant discarded. The pellet was resuspended in 200 μl of RNase A/PBS (100μg/ml) and incubated for 2 h at 37°C in the dark. Propidium iodide (Fisher Scientific, P3566) was added at a final concentration of 5μg/ml just before analysing the samples by flow cytometry using the ACEA Novocyte flow cytometer. The analysis was performed using the NovoExpress® software.

### Microccocal nuclease treatment

HeLa cells were transfected with 1 μg of GFP, GFP:H2A.Z.2.1^WT^ or H2A.Z.2.1^KR^ for 24 h. Cells were lysed using lysis buffer (1M Tris-HCl, 2.5M NaCl, 10% NP-40) and the chromatin was extracted by centrifugation at 1000 g for 3 min. Chromatin was flash frozen in liquid nitrogen until ready to use. Chromatin was digested with Microccocal Nuclease (NEB, 37°C, 30 min) in digestion buffer (1M Tris-HC, 1M CaCl_2_, 0.5M MgCl_2_, 2M sucrose).

Digested chromatin was resuspended in Laemmli sample buffer and run in a 10% acrylamide gel. Anti-GFP antibody (Roche, Cat#11814460001) was used to detect the construct and the anti-H3 C terminus (Active Motif, Cat#39052) as a control.

To verify that chromatin was digested, 1% of SDS (final concentration) was added to 50 μl of digested and undigested chromatin. DNA was extracted with phenol chloroform, precipitated with ethanol and analysed in a 1% agarose gel.

### RNA sequencing

RNA was collected from siRNA-treated HeLa cells and extracted using the RNeasy PowerLyzer Tissue & Cells Kit (Qiagen) according to manufacturer’s protocol. RNA samples were sent to the Wellcome Trust Genomic Centre (Oxford University, Oxford UK) for whole-genome sequencing, using the Illumina HiSeq4000.

The RNA-seq data was analysed using open source software from the Tuxedo suite, including TopHat2 ^87^ and Cufflinks ^88^. The paired end raw reads were mapped to the human reference genome GRCh38 using the annotations from GENCODE 28 ^89^, with TopHat2 v2.1.1 (Bowtie 2 v2.2.6) under standard conditions. All the sequence and annotation files were downloaded from the Illumina website. The resulting alignments were filtered for high quality hits using Samtools v0.1.19 ^90^ with a minimum selection threshold score of 30. Cufflinks v2.2.1 was used to assemble the mapped reads into transcripts and quantify their expression levels. Finally, Cuffdiff, part of the Cufflinks package, was used to identify differentially transcribed genes between samples. Functional enrichment was analysed using String (string-db.org), while Venn diagrams were performed in the open software FunRich. Volcano plots were performed using the ggplot package in R v3.5.0.

The file containing the H2A.Z ChIP-seq data used (E117-H2A.Z.narrowPeak.gz. https://egg2.wustl.edu/roadmap/data/byFileType/peaks/consolidated/narrowPeak/) were filtered using a q-value threshold of 0.01. The transcription start sites (TSS) are defined by Abugessaisa et al. ^91^.

If two or more H2A.Z regions overlap, they were considered as one region. For each set of genes, the H2A.Z regions that were not within +/-5 kb of any of the genes in that set were excluded, and the remainder were classified into 3 categories: regions that overlap and/or are within +/-3 kb of one or more of the TSS associated with a gene whose expression changes after RNAi knockdown; regions that overlap the loci of genes whose expression changes after RNAi knockdown but not any of the TSS associated with those genes; and regions that do not overlap the TSS nor the loci of genes whose expression changed after RNAi knockdown. The number of H2A.Z regions in each category is divided by the total number of regions for that gene set to give the proportion of H2A.Z regions in each category.

### Differentially expressed genes frequency analyses

To calculate the expected and observed frequency of differentially expressed genes on each chromosome, we used the gene numbers present on each human chromosome and used the chromosome copy number in HeLa cells integrating both the published datasets (10.1371/journal.pone.0029225) and our own experimental dataset for chromosomes 17 (this study) and chromosomes 13 and 14 ^31^.

### Statistical analyses

Statistical analyses were performed either in Excel (Chi-square test), or in R (using the Wilcoxon rank test function, differential expression, lowess smoothing).

## SUPPLEMENTARY FIGURE LEGENDS

**Supplementary Figure 1.**
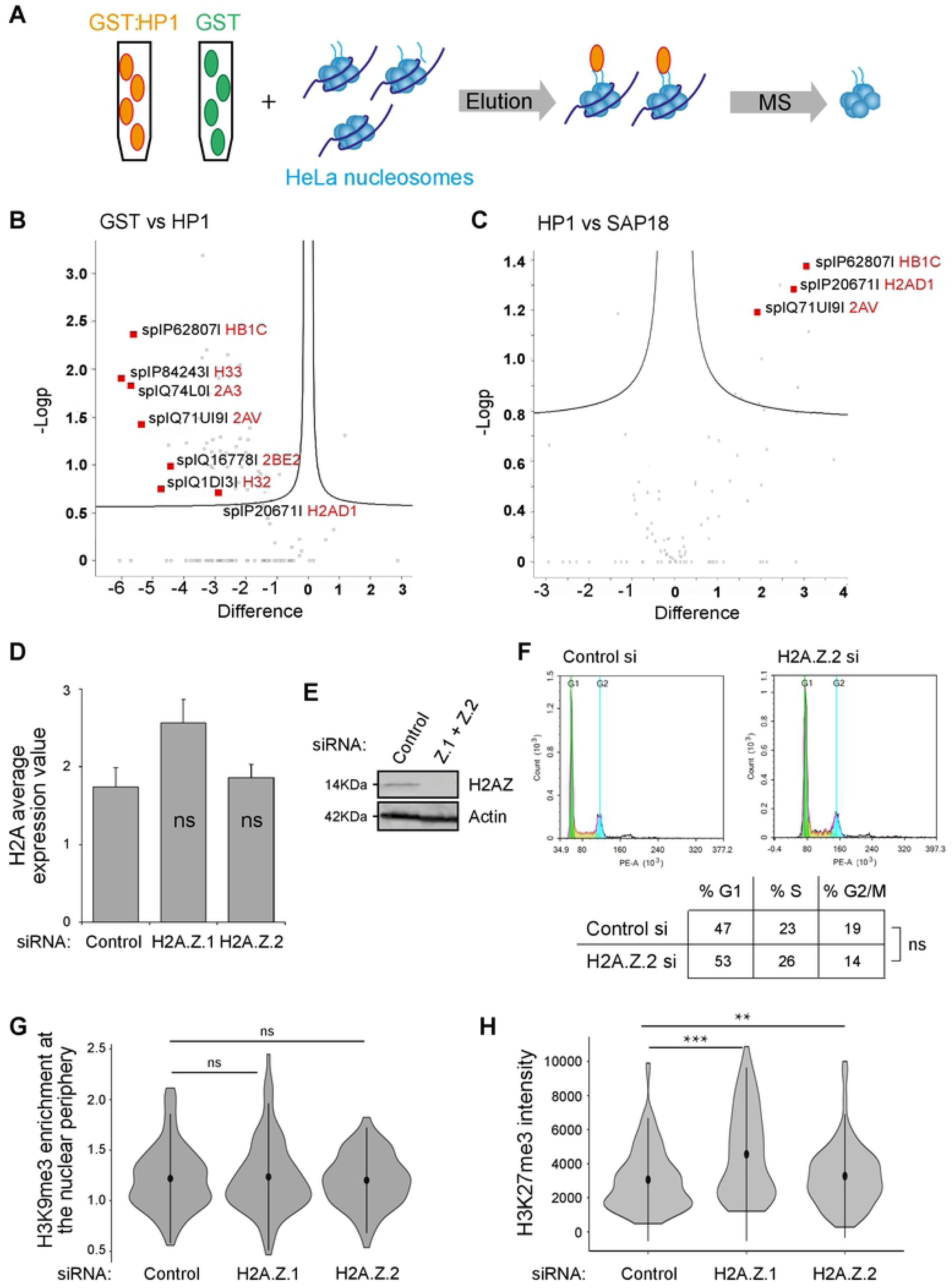
H2A.Z.2 is necessary for heterochromatin maintenance in human cells. **A)** Scheme of the proteogenomic approach. **B)** Volcano plot showing the enrichment of histone variants in the GST fraction compared to HP1. **C)** Volcano plot showing the enrichment of histone variants in the HP1 fraction compared to SAP18. **D)** *H2A* average expression values obtained by RNA sequencing of three biological replicates after control H2A.Z.1 or H2A.Z.2 siRNA treatment. Error bars show the standard deviation (SD). ns=not significant. **E)** Western blot of Control and H2A.Z.1 + H2A.Z.2 RNAi. Whole cell lysates were loaded and probed with anti H2A.Z or beta actin antibodies **F)** Cell cycle profile by FACS of control and H2A.Z.2 siRNA-treated HeLa cells. Percentages represent the mean of two biological replicates. Data sets were statistically analysed using Chi-square test. ns=not significant. **G)** Violin plots of the enrichment of H3K9me3 at the nuclear periphery in cells from the experiment in figure 1 D. Mean and SD are shown. Data sets were statistically analysed using the Wilcoxon rank test in R. ns=not significant. **H)** HeLa GFP:HP1a were transfected with control, H2A.Z.1 and H2A.Z.2 siRNAs, fixed, and stained for H3K27me3. Violin plots represent the distribution of H3K27me3 intensity levels of at least 200 nuclei from four biological replicates. Mean and SD are shown. Data sets were statistically analysed using Wilcoxon rank test. **=p<0.01; ***=p<0.001.

**Supplementary Figure 2.**
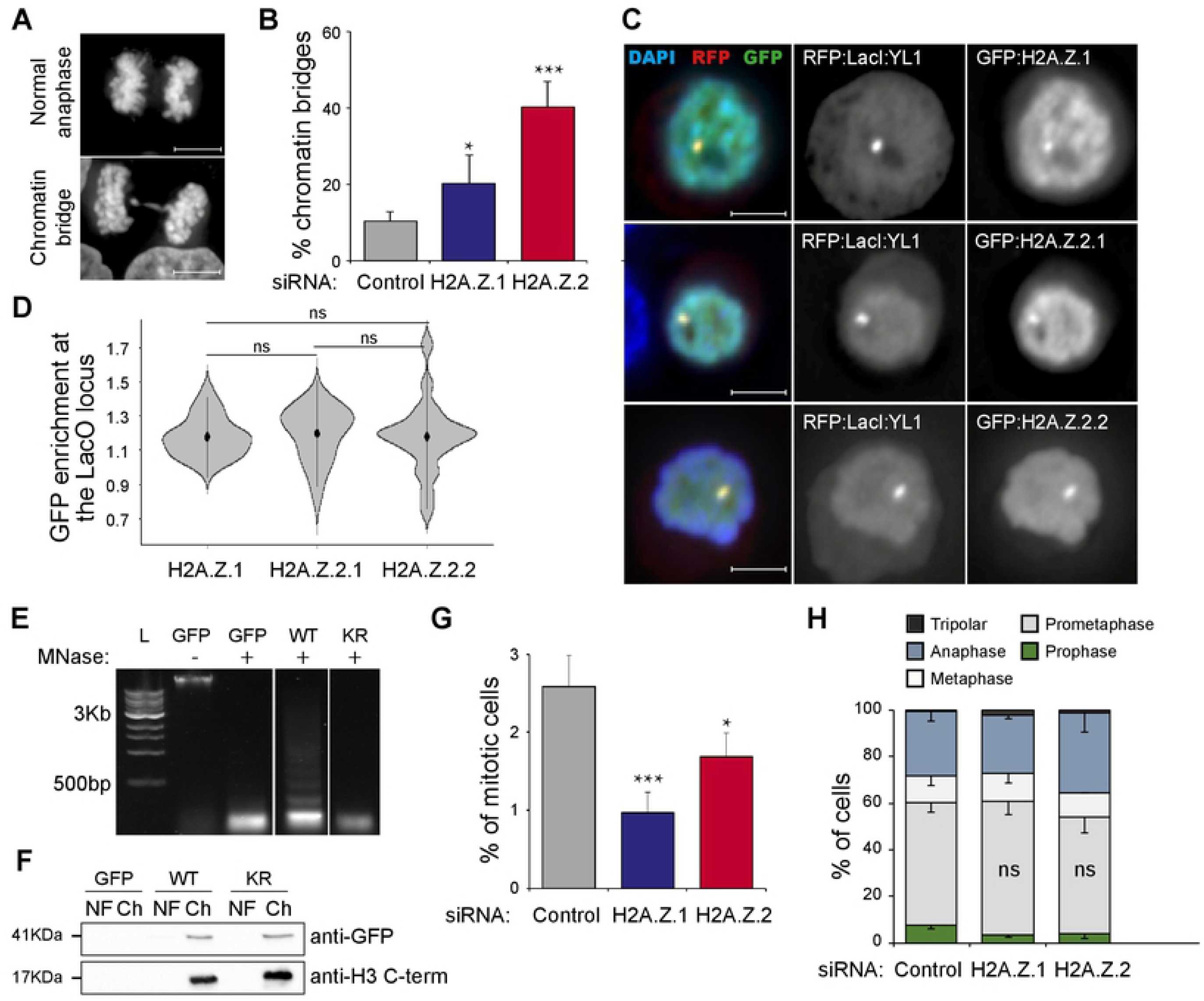
H2A.Z.2.1 knockdown leads to genome instability. A) Example of a normal anaphase (top panel) and an anaphase with a chromatin bridge (bottom panel). Scale bar: 10 μm. B) Quantification of the number of anaphases with chromatin bridges of HeLa cells after transfection with control, H2A.Z.1 or H2A.Z.2 siRNA. Error bars represent SD of three biological replicates. At least 100 anaphases were analysed for each condition. Data sets were statistically analysed using Chi square test. *=p<0.05; ***=p<0.001. C) Representative images of DT40 cells carrying a LacO array inserted at a single locus co-transfected with RFP:LacI:YL1 (red) and either GFP:H2A.Z.1, GFP:H2A.Z.2.1 or GFP:H2A.Z.2.2 (green). Scale bar: 5 μm. D) GFP enrichment was calculated as a ratio between the intensity at LacI spot and the mean of two random nuclear spots. Mean and SD are shown. Data sets were statistically analysed using Wilcoxon rank test. ns=not significant. E) HeLa cells were transfected with GFP, GFP:H2A.Z.2.1WT (WT) or GFP:H2A.Z.2.1KR (KR), lysed, and digested with micrococcal nuclease (MNase) for 30 min to generate mononucleosomes. L: DNA ladder. F) The chromatin fraction (Ch) from E was separated on SDS PAGE gel, together with the nuclear fraction (NF), and subjected to GFP immunoblotting. Anti-H3 C terminus antibody was used as a control. G) The percentage of mitotic cells was calculated from HeLa cells transfected with control, H2A.Z.1 or H2A.Z.2 siRNA. Error bars represent SD of three biological replicates. At least 2500 cells were analysed for each condition. Data sets were statistically analysed using Chi-square test. *=p<0.05. H) Mitotic cells from the experiment in G were analysed and classified by mitotic stage. Error bars represent SD of three biological replicates. At least 300 mitotic cells were analysed for each condition. ns=not significant.

**Supplementary Figure 3.**
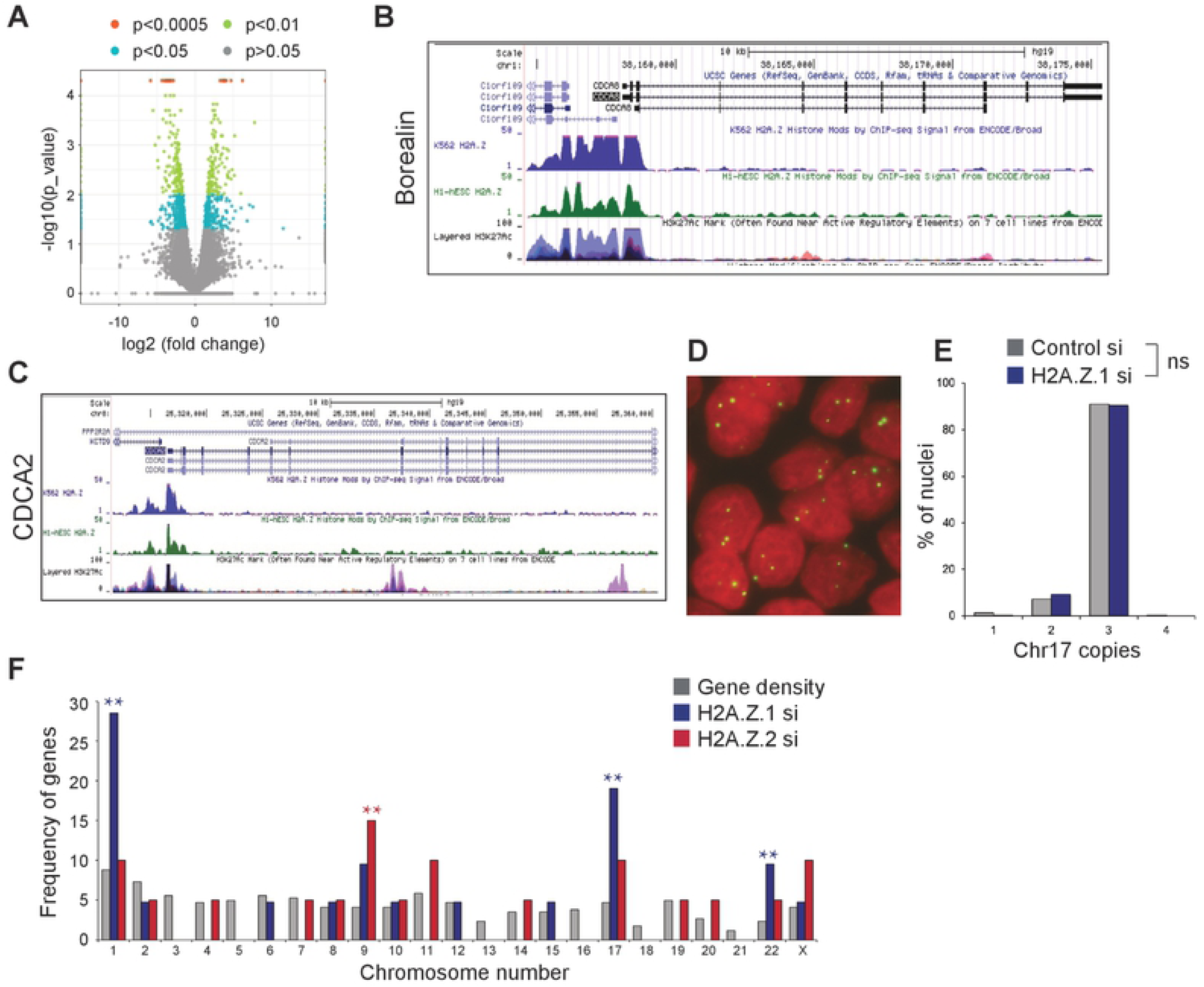
H2A.Z.1 regulates cell cycle progression. **A)** Volcano plot representation of differentially expressed genes in the H2A.Z.1-depleted cells vs the H2A.Z.2-depleted cells data sets. Y axis represent the -log10 of the p value. Colour points mark the genes with significantly increased or decreased expression. X axis represent the log2 value of the fold change: points with log2<0 indicate downregulated genes in the H2A.Z.2-depleted cells data set compared to the H2A.Z.1-depleted cells data set; log2>0 indicate upregulated genes. **B)** UCSC analyses of H2A.Z localisation on Borealin showing H2A.Z enrichment at the TSS. **C)** UCSC analyses of H2A.Z localisation on CDCA2 showing H2A.Z enrichment at the TSS. **D)** Representative image of HeLa nuclei after FISH with a Chr17 centromeric probe (green). **E)** Quantification of the number of FISH signals/nucleus in control (grey) or H2A.Z.1-depleted (blue) cells. At least 500 nuclei were analysed per condition. Data sets were statistically analysed using Fisher exact test. ns=not significant. **F)** Frequency of expected (grey) vs observed genes per chromosome after H2A.Z.1 (blue) or H2A.Z.2 (red) depletion. Data sets were statistically analysed using Fisher exact test. **=p<0.01.

## ACKNOWLEDGMENTS

We thank Prof Sala’s group (Brunel University London) and Prof Wendy Bickmore (Edinburgh) for advice and useful discussions.

The Vagnarelli lab is supported by the Wellcome Trust Investigator award 210742/Z/18/Z to Paola Vagnarelli.

This work was partially supported by the Brunel Idea Award to PV and IJDC.

RSG was a recipient of the Isambard PhD scholarship (2016-2019) and now is supported by the Wellcome Trust.

The Wellcome Centre for Cell Biology is supported by core funding from the Wellcome Trust [203149].

## AUTHOR CONTRIBUTIONS

Conceptualization, P.V., R.S.G.; Methodology, I.J.DC., C.S., J.R.; Investigation, R.S.G., I.J.DC., H.A, C.S, C.S and P.V.; Writing – Original Draft, P.V. and R.S.G.; Writing – Review & Editing, all the Authors; Funding Acquisition, P.V., I.J.DC and JR; Supervision, P.V., C.S., V.V.

## DATA AND MATERIALS AVAILABILITY

All unique/stable reagents generated in this study are available from the Lead Contact with a completed Materials Transfer Agreement.

The Mass spectrometry datasets generated during this study are available via ProteomeXchange with identifier PXD018719

Reviewer account details:

**Password:** cBIH10fL

The RNA seq datasets generated during this study are available at NCBI http://www.ncbi.nlm.nih.gov/bioproject/629054 (SubmissionID: SUB7307224).

The microscopy images supporting the current study are available from the corresponding author on request and they will be shared via FIGSHARE.

